# Spiral-Sensing and Fold-Change Detection Direct Ascidian Sperm to the Egg

**DOI:** 10.1101/2025.09.16.676465

**Authors:** Makoto Iima, Kogiku Shiba, Manabu Yoshida, Kazuo Inaba, Toshiyuki Nakagaki

## Abstract

From microorganisms to marine sperm, successful navigation toward chemical cues relies on the integration of sensory inputs with locomotion. Ascidian sperms exhibit a robust chemotactic strategy by swimming along circular paths whose centres form large-scale spirals, guiding them toward attractant-releasing eggs. Although their intracellular Ca^2+^ dynamics exhibit rhythmic bursts aligned with flagellar modulation, the underlying computational principles remain unclear. Here, we propose a minimal model that captures ascidian sperm chemotaxis through a combination of swimming trajectory control and an internal signalling model with fold-change detection (FCD). This model responds to relative rather than absolute changes in the stimulus concentration, achieving scale-invariant signal processing. Concurrently, the spiral swimming path enables spatial sampling of the concentration field, allowing the cell to infer gradient direction. Our findings demonstrate that the integration of a resilient signalling module and physically structured sampling trajectory enables effective chemotaxis in complex environments.

## 1 Introduction

Animal locomotion plays an essential role in fertilisation, mating, foraging, and predator avoidance. To maximise survival and reproductive success, animals have evolved diverse strategies such as swimming, flying, walking, and crawling adapted to their environments. Locomotion typically involves three integrated components, namely kinematics, sensory detection, and signal processing. Understanding such tightly coupled systems is the key to uncovering the principles of biological movement. Simplified organisms, including protists and motile cells that exhibit chemotaxis, provide tractable models for studying environmentally guided navigation systems.

This study focused on chemotaxis and the ability to detect and respond to chemical gradients. Chemotaxis is a ubiquitous phenomenon observed in diverse organisms ranging from bacteria to amoebae [1, 2, 3, 4], motile cells such as neutrophils and sperm [5, 6, 7], and even inanimate objects [8, 9] .

The chemotaxis mechanism of *Escherichia coli* is based on the regulation of multiple flagella driven by rotary motors. Alternating counterclockwise (CCW) rotation provides linear movements (“run”); clockwise (CW) rotation causes tumbling movements (“tumble”) for directional changes. Transitions between these states occur approximately every second under steady-state conditions [10].

In a chemical gradient environment, *E. coli* adapts by extending runs and reducing tumbles towards higher attractant concentrations [10]. Mathematical models, such as those developed by Tu et al. [11], have elucidated the underlying signalling pathways and fold-change detection (FCD) properties, a property of biological sensory systems wherein where the output signal remains unchanged in form, even when the input ligand concentration increases by a constant factor [10]. FCD has been reported in various biological sensory systems; chemotaxis of *E. Coli* [12], cAMP response of social amoebae[13], circadian clock of murine fibroblasts[14], visual processing of *Ciona* larvae[15], to name a few. Sperm chemotaxis is crucial for reproduction because it guides sperms to eggs via attractant detection. Unlike *E. coli*, sperm use a larger eukaryotic flagellum; their typical circular or helical swimming patterns reflect distinct chemotactic mechanisms [16].

In sea urchins, chemotaxis involves flagellar receptors, intracellular signalling, Ca^2+^ modulation, altering flagellar dynamics and trajectories [17, 18] Ascidians show similar process but with specific differences in receptor types and signalling mechanisms [19, 20].

Shiba et al. [21] investigated Ca^2+^ dynamics and trajectories in a gradient of the attractant concentration — sperm-activating and attracting factor (SAAF) [22]. They found that Ca^2+^ bursts occurred at local SAAF minima along the orbit, increasing the curvature and initiating straight swimming before returning to a circular trajectory. This dynamic adjustment mechanism enables their navigation of the attractant.

Mathematical models of sperm chemotaxis have elucidated key links among cell trajectories, signal transduction, and environmental cues [23, 24, 25, 26]. Friedrich and Jülicher [6, 27] proposed a general framework representing chemotactic concentration fields as *c*(*r*), with the stimulus *s*(*t*) = *c*(*r*(*t*)), an output *a*(*t*), and an internal variable *p*(*t*) for signal processing. Although the model captures steady-state adaptation, it lacks the FCD property.

Here, we present a model specifically designed for ascidian sperm chemotaxis that incorporates explicit biochemical signalling dynamics. The model reproduced key experimental observations, including a consistent phase shift between internal signalling and SAAF, as well as characteristic swimming trajectories in both two- and three-dimensional spaces. Moreover, it exhibited FCD, which was evaluated in light of the first experimental evidence of FCD in ascidian sperm. This framework enhances the mechanistic understanding of chemotaxis and provides a foundation for developing biomimetic systems with navigational functionality.

## 2 Results

### 2.1 Signal transduction

Figure 1A illustrates a conceptual diagram of the signal transduction model (for the model details, see Methods and SI2). We first analysed the relationship between the attractant (SAAF) concentration, *s* (as the input signal), and the intracellular Ca^2+^ concentration, *c* (as the output) by examining Eqs. (3) and (4), respectively. Since plasma membrane Ca^2+^-ATPase (PMCA) has been proposed as a receptor for SAAF, the present model established a circuit in which SAAF reception and Ca^2+^ efflux are directly linked. This system will hereafter be referred to as the “signal transduction system.”

**Figure 1:**
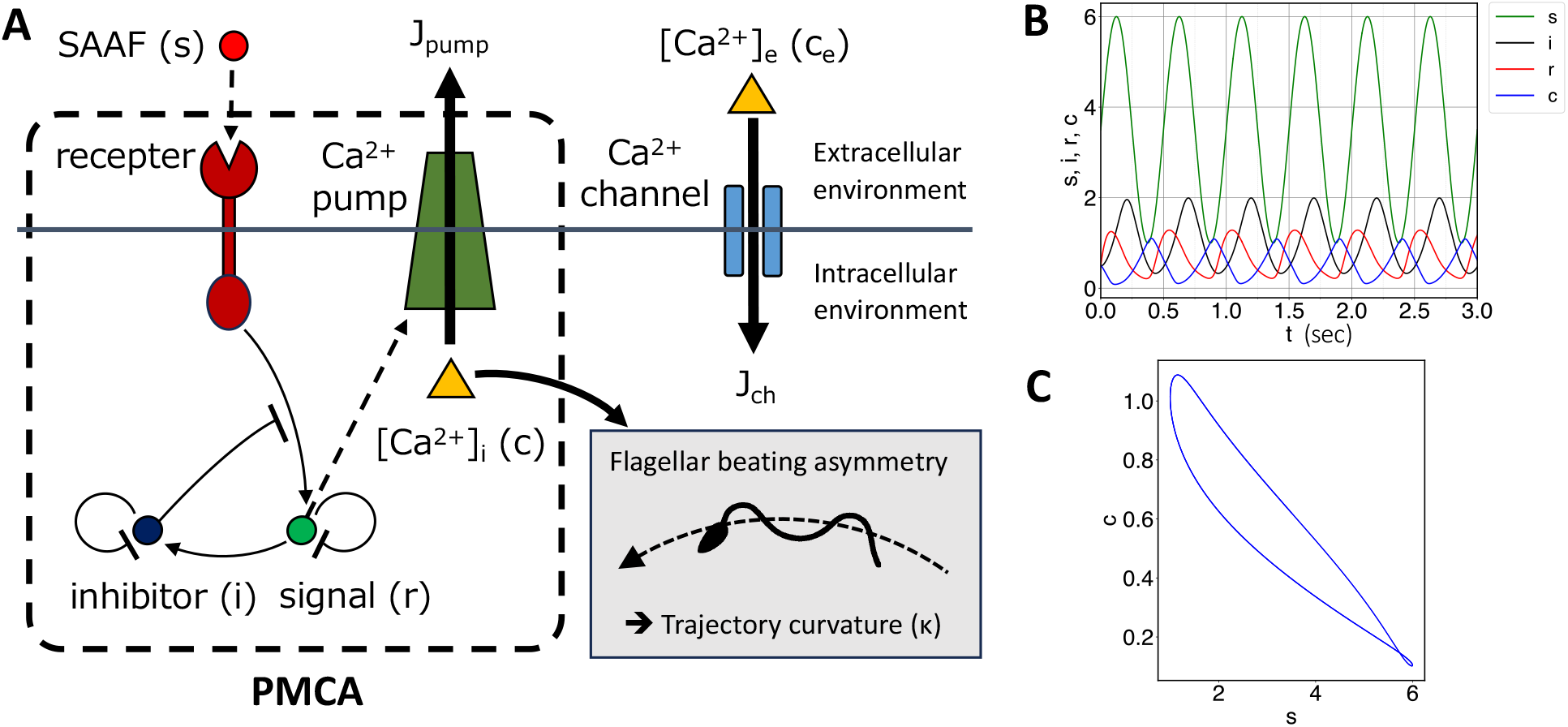
A: Conceptual diagram of the signal transduction model. The functions of PMCA are incorporated into the signalling module and the Ca^2+^ pump. The SAAF signal, *s*(*t*), regulates the Ca^2+^ pump via a feedback control circuit with fold-change detection (FCD) properties. The intracellular Ca^2+^ concentration, [Ca^2+^]_*i*_, is dynamically modulated by the Ca^2+^ pump and a Ca^2+^ channel, which is influenced by the extracellular Ca^2+^ concentration. B: Time series of the system response to a sinusoidal input signal, *s*(*t*), where *s*_min_ = 1. C: Trajectories in the phase space (*s*(*t*), *c*(*t*)) during the latter half of the simulation, where *c*(*t*) represents the intracellular Ca^2+^ concentration.

The input signal *s* is modelled as a sinusoidal function that mimics the signal experienced by the sperm swimming along a circular trajectory in a linear gradient of the attractant: 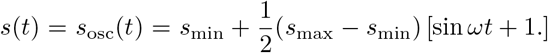 The parameters and simulation conditions are provided in SI 3. A typical case of the time series of *s* = *s*_osc_, *r, i*, and *c* is shown in Fig. 1B. The variables *r, i*, and *c* converge to periodic functions within one period. Notably, when *r* peaks, *c* takes smaller values, and *c* rises as *r* decreases. The peak of *c* aligns closely with the minimum of *s*.

This result is consistent with experimental observations indicating an increase in the intracellular Ca^2+^ concentration near the SAAF minima (distal phase) along the sperm trajectories [21]. The antiphase relationship between *s* and *c* is further illustrated in the phase space (*s, c*), as shown in Fig. 1C.

The signal transduction system exhibits an extended fold-change detection (FCD) property when *s* is treated as the input, *i* is the internal variable, and *r* and *c* are the outputs [10] (SI 4). Notably, the antiphase relationship remains robust over a wide range of *s*_min_, *s*_max_, and input signal periods (SI 5).

### 2.2 Chemotactic trajectory to a point source

We calculated the chemotactic systems described by Eqs. (3), (4), and (5) in the presence of an *s* field (see SI 2, 3). This system also demonstrated an extended fold-change detection (FCD) property (SI 4).

First, we considered the case where the *s* field is defined as:

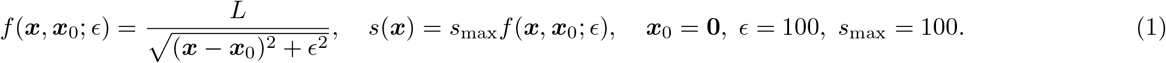

Here, *s* is expressed in nanomolar units of the same order of experimental values and *L* is a length scale. This setup simulated a scenario in which a chemotactic substance was localised at a point source, creating a concentration gradient through diffusion. The steady-state solution of the diffusion equation from a point source in three dimensions is proportional to |*x* −*x*_0_| ^−1^ for large |*x* − *x*_0_|, consistent with the asymptotic behaviour of Eq. (1). We assumed that *s*(***x***) was unchanged by the sperm motion.

The trajectory of the sperm model (treated as a point) in physical space is shown in Fig. 2A. Initially positioned on the right (denoted by a red point, with the initial direction indicated by an arrow), the sperm model followed a helical trajectory, ascending the *s* concentration gradient until it reached a region near the peak of *s*. This trajectory qualitatively resembled the experimental observations, particularly in the distal phase, where the curvature increases [21]. Notably, this behaviour is reproduced without explicitly introducing the time derivative of the SAAF or a finite time delay as used in other models [23, 24]. Furthermore, the model captures the chemotactic behaviour without incorporating a detailed response mechanism for curvature changes [21, 25].

**Figure 2:**
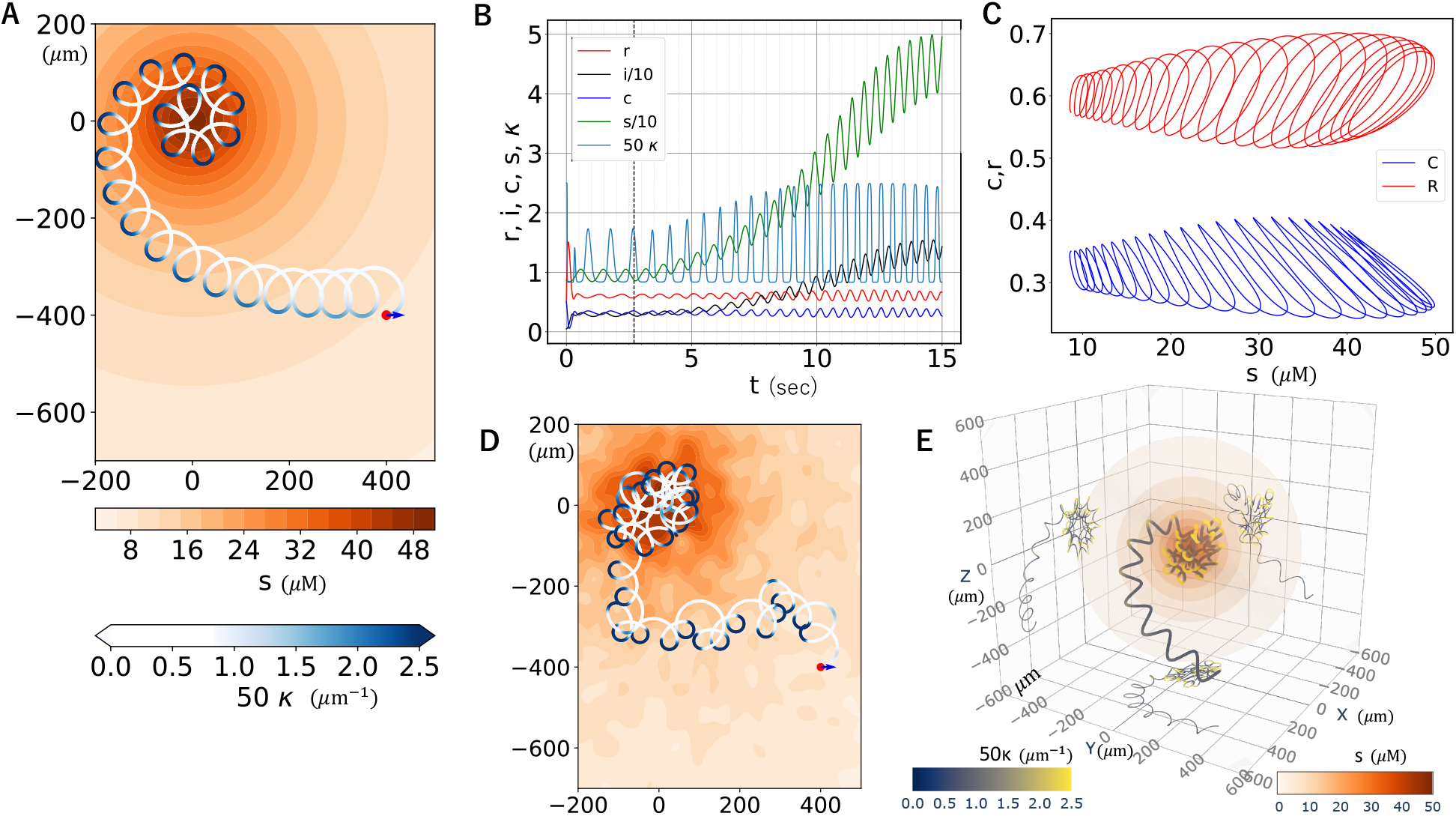
A: Trajectories of the sperm model in the physical space within a SAAF (*s*) field decaying from a point source. The background colour represents the concentration of *s*, whereas the trajectory colour corresponds to the curvature. B: Time series of the signal transduction variables, *s, i, r, c*, and the trajectory curvature *κ*. C: Trajectories in the phase space, (*s, r*) and (*s, c*). D: Trajectory of the sperm model under the perturbed SAAF field with relative amplitude *α*(= 0.1) and the correlation length *l*(= 25 µm) (2.7 *< t <* 20). E: Trajectory of the 3D sperm model for 2.7 *< t <* 20, with a torsion value of 0.01. Spheres indicate isosurfaces of *s*−field, while the trajectory colour denotes the curvature. Projections onto the *xy*−, *yz*−, and *zx*−planes are also shown.

A time series of this process is shown in Fig. 2B. Along the trajectory, value of *s* has an increasing trend. The curvature radius during this period ranges approximately from 70 to 100*µ*m, values that closely match experimental measurements [21]. The change in *c* while ascending the gradient corresponds to the orbital curvature *κ* as defined, with the peak times of *c* and *κ* coinciding. Additionally, the minimum of *s* roughly aligns with the maximum of *κ*, illustrating an antiphase relationship between *s* and *c*.

This antiphase relationship is further illustrated by the (*s, c*) trajectory shown in Fig. 2C. Conversely, *r* was in phase with *s*, indicating a time delay in the dynamics of intracellular Ca^2+^. These properties are consistent with the signal transduction model, in which *s* is represented by a sinusoidal input. The robustness of signal transduction for varying *s*_min_ and *s*_max_ values (see SI 5) reflects the robustness of the chemotaxis model.

To examine a more realistic scenario, we analysed the trajectory under a perturbed SAAF field generated by a Gaussian process (using a Gaussian kernel with correlation length *l* µm; see SI 3 for details), in order to mimic the mixing effects caused by other sperms and fluid flows. Figure 2D presents a representative example for the case *l* = 25 µm, which is of the same order as the trajectory radius, and *A*, the relative amplitude of the noise, is set to 0.1 (Statistical analysis is in SI 6). In the three-dimensional sperm model (see SI 3), a typical trajectory is shown in Fig. 2E. Even under a random initial orientation, the model achieved a success rate of 67%, despite some trajectories initially moving away from the source (SI 6). In both scenarios, the sperm model successfully navigated towards a region near the concentration peak, demonstrating distinct aspects of its robustness.

Finally, to confirm the FCD property in ascidian sperm, we experimentally investigated how the trajectory characteristics change under varying background SAAF concentrations. In the experimental setup shown in Fig. 3Aa, the SAAF concentrations at the source is *b µ*M and the background field consisted of artificial seawater (ASW) with SAAF concentrations of *a* nM, because a certain SAAF concentration was necessary to activate the sperm. At every point in space and at every moment in time from the start of the experiment, these SAAF fields exhibit a proportional relationship, satisfying a condition of input signal for fold-change relationship. Figure 3B shows typical trajectories for (*a, b*) = (1, 1), (2, 2) and (5, 5). Despite the strength of SAAF field for (*a, b*) = (5, 5) is five times of that for the (*a, b*) = (1, 1) at every point in space, the swimming trajectory toward the source showed marked similarity. The parameters used to evaluate chemotactic turns such as radial velocity magnitude (Fig. 3Ab; the definitions is in SI 7), period, displacement between turns and maximum and minimum curvatures of the trajectories were assessed (Fig. 3C and SI 7). These also did not exhibit statistically significant responses, except for the period and maximum curvature at (*a, b*) = (1, 1). Considering that the initial conditions of the orbits were not the same, these results suggest that FCD holds true in ascidian sperm trajectories in this range of SAAF concentrations.

**Figure 3:**
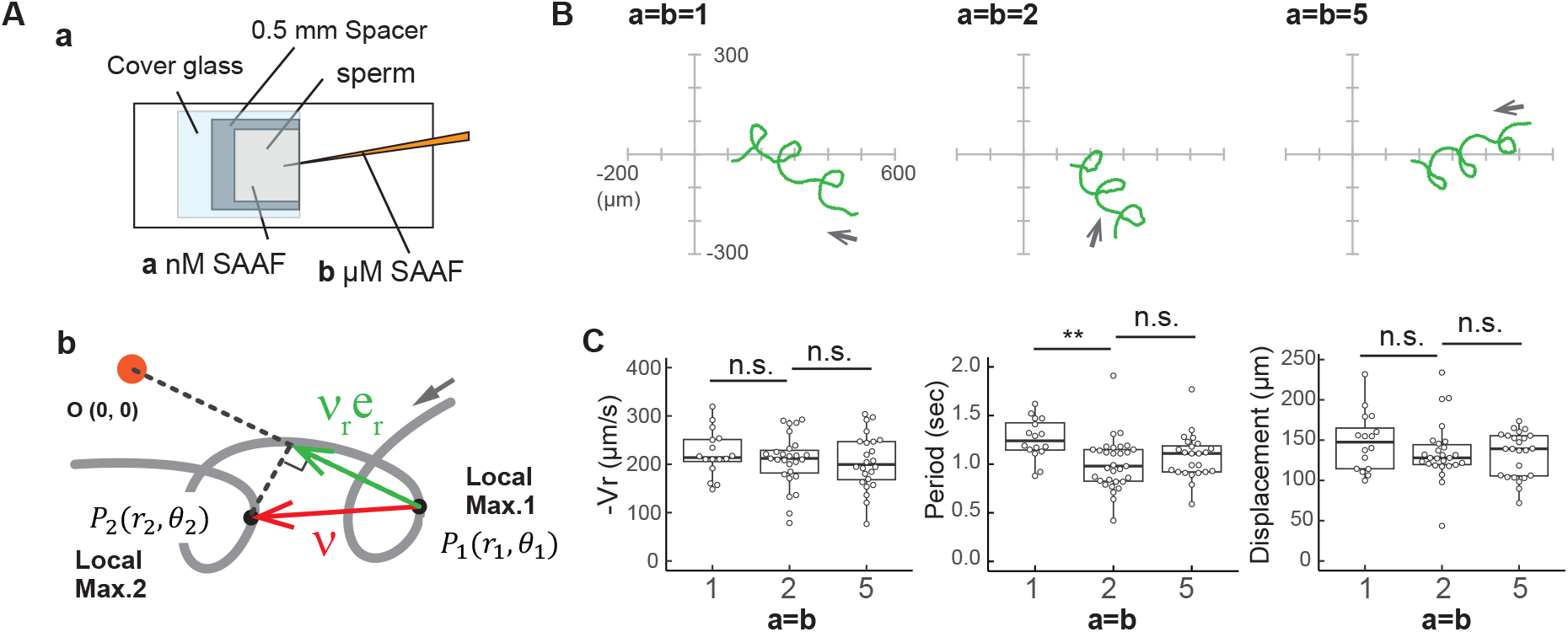
A: Experiment of FCD property demonstrated by ascidian sperm under three conditions: (*a, b*) = (1, 1), (2, 2) and (5, 5), where the SAAF concentration at the source is *b µ*M diluted in ASW with 1% agar and the SAAF concentration in the background field is *a* nM diluted in ASW. a: The observation chamber for analysis in a SAAF concentration field from a point source. b: Schematic figure of swimming trajectory toward the chemoattractant source. Arrow indicates sperm swimming direction. Times giving local maxima distance from the source are used to numbering the turns. The mean velocity between turns, ***v***, is defined by the displacement and time difference between adjacent turns. B: Typical sperm swimming trajectories. Arrow indicates sperm swimming direction. C: The radial component of *v*, −*v*_*r*_, the period and the displacement between turns when the sperm approaches the source. Sample numbers *N* = 16 (1-1), 31 (2-2) and 24 (5-5) from three different experiments. Distribution of values is plotted in a box plot. ** Significant at *p <* 0.01(Dunnett test), as compared with the value of (*a, b*) = (2, 2).

### 2.3 Trajectories under one-dimensional attractant gradient and a canonical trajectory

A one-dimensional SAAF gradient was considered. This situation is useful for the definition of a canonical trajectory of this model, by which the quantitative chemotactic performance assessment is possible, and corresponds to experimental conditions where the SAAF field diffuses from a linear distribution of the source.

First, we considered a linear SAAF field, 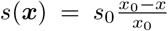, where *s*_0_ = 30 µM and *x*_0_ = 600 µm. The resulting trajectory is as shown in Fig. 4A. The displacement between turns progressed at an angle to the gradient of *s*(*x*), and chemotactic behaviour was observed under these conditions. This angle gradually decreases as the gradient of the sperm model increases. The curvature *κ* reaches its maximum near the distal phase, where the SAAF concentration is minimal, consistent with experimental observations [21].

**Figure 4:**
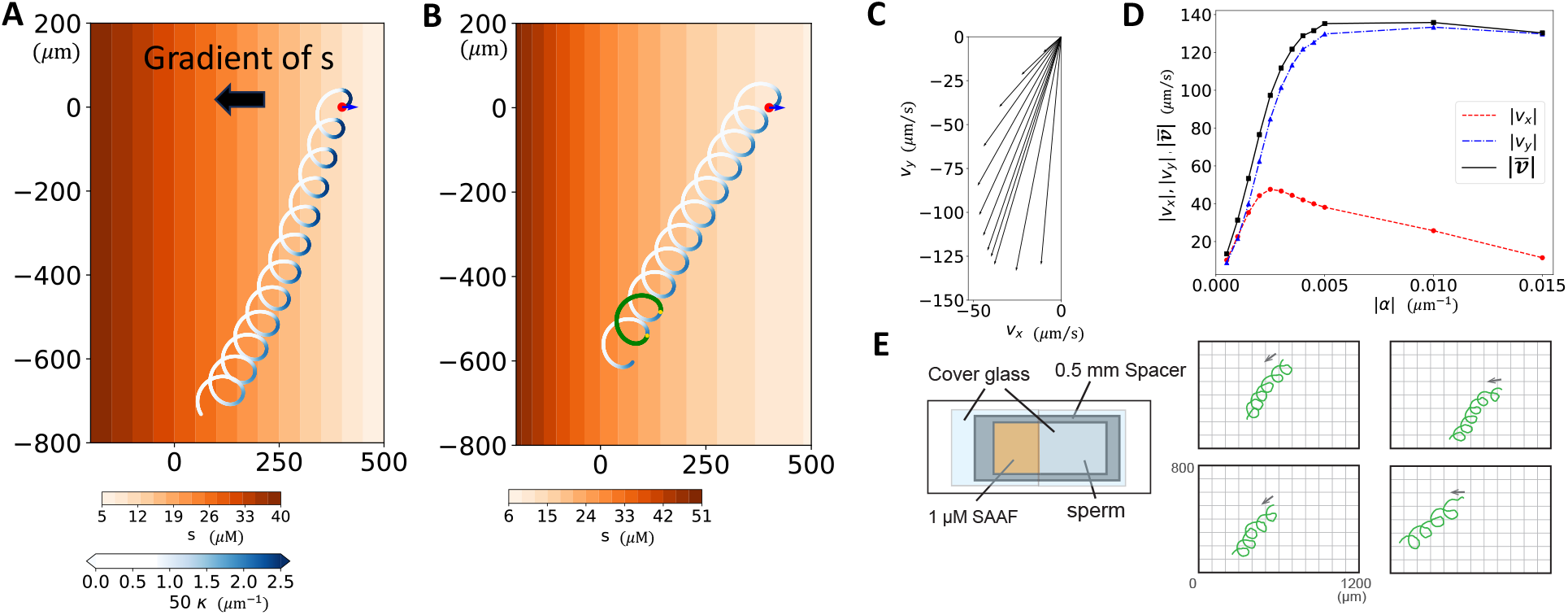
(A) Trajectories of the sperm model in physical space within a 1D linearly distributed SAAF field. The view of the trajectories is the same as in Fig. 2A. (B) Trajectories of the sperm model in physical space within a 1D exponentially distributed SAAF field. The scale bar of *κ* is the same as Fig. 4A. Two yellow points represent consecutive maxima of *κ*, and the trajectory advanced during one cycle is highlighted in green. (C) Average velocity vectors of the sperm model for different decay rates *α* of the exponential distribution. (D) Average velocity vectors for various values of *α*, including plots of the average magnitude of the velocity vector ***v*** and its components *v*_*x*_ and *v*_*y*_. (E) Experimental trajectories of ascidian sperm. The observation chamber and sperm swimming trajectories in a one-dimensional distribution of SAAF field. Arrow indicates sperm swimming direction.

It is noteworthy that the distance travelled per period, defined as the difference between consecutive times when *κ* reaches its maximum, gradually decreases as the sperm ascends. This behaviour occurs because the relative increase in the SAAF concentration during a period diminishes in the linear distribution, making this distribution less suitable for characterising the trajectory.

To better characterise the trajectory under a one-dimensional concentration gradient, we considered an exponential distribution:

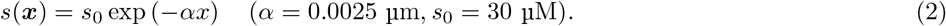

In this case, the spatial decay rate, −(∂*s/*∂*x*)*/s*, is a constant, *α*. The trajectory of this distribution is shown in Fig. 4B. Here, the trajectory exhibited a constant increment vector per period.

Notably, the mean velocity over one period 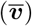, increment length divided by the period, is independent of the sperm position, which can be proved based on the FCD property (SI 6). In Fig. 4B, the trajectory during one cycle is highlighted in green. Fig. 4C,D illustrate the average velocity 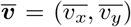 for different decay rates *α*. The average velocities 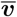 for various *α* values are shown as arrows in Fig. 4C, where *α* = 5.0 *×* 10^−4^*j* (*j* = 1, 2, …, 12) from top to bottom. Fig. 4D presents |*v*_*x*_| and |*v*_*y*_|, as well as the speed 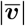. The value of 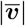 reaches its maximum at *α* = 0.005, whereas the velocity component in the gradient direction, 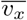, peaks at *α* = 0.0025.

The corresponding experimental results are presented in Fig. 4E (see also SI 1). A one-dimensional concentration gradient was generated by using agar soaked in SAAF. The resulting distribution, governed by the one-dimensional diffusion equation, is time-dependent and does not exhibit a linear shape, depicting a concavity similar to that of an exponential distribution.

The concentration gradient was predominantly horizontal; however the mean trajectory formed an angle of approximately 45^*°*^ with respect to the gradient direction. The curvature increased in the distal phase, and the observed reduction in the mean velocity as the concentration increased aligned with the model trajectory. These findings suggest good agreement between the experimental and model trajectories.

### 2.4 Chemotaxis property dependency on efflux

Ca^2+^ influx through Ca^2+^ channels is controlled by the parameter *j*_0_ in the model. Figure 5A shows the period-averaged velocities in an exponential SAAF field (Eq. (2)) for various values of *j*_0_ (see Methods; Eq.(4)), which ranges from 0.50 to 0.65 in increments of 0.01. The magnitude of the period-averaged velocity, 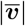, reached a maximum at *j*_0_ = 0.63, whereas the magnitude of the velocity component in the gradient direction, |*v*_*x*_|, peaks at *j*_0_ = 0.60. We selected three representative cases: (a) *j*_0_ = 0.55, (b) *j*_0_ = 0.60, and (c) *j*_0_ = 0.65. The trajectory for case (b) is shown in Fig. 4B, where significant chemotactic behaviour is observed. In case (a), low Ca^2+^ influx resulted in a limited increase in the intracellular Ca^2+^ concentration, leading to weak chemotaxis (Fig. 5Aa). Conversely, in case (c), the high Ca^2+^ influx maintained a high intracellular Ca^2+^ concentration, also resulting in weak chemotaxis (Fig. 5Ac).

**Figure 5:**
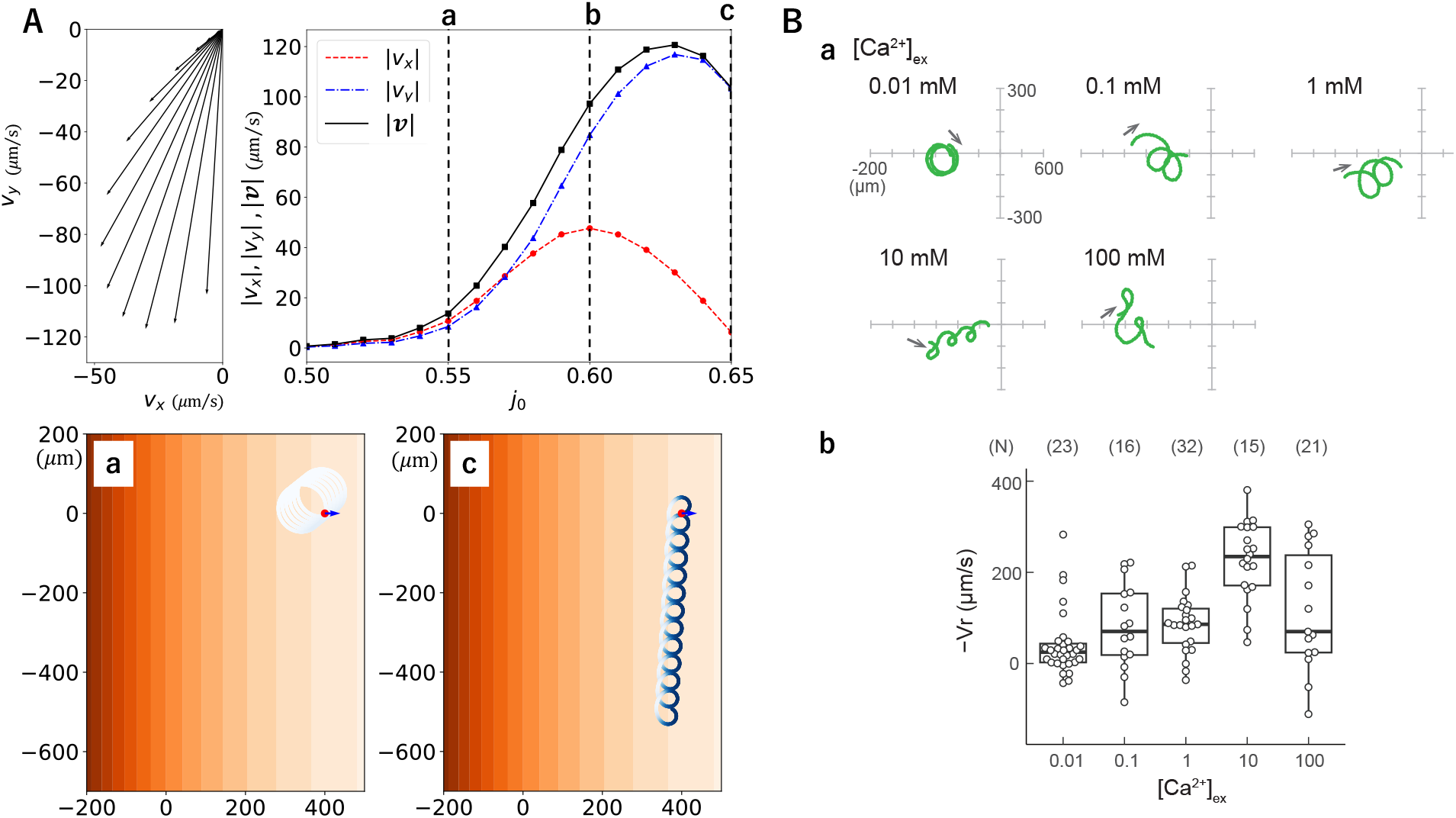
A: Characterization of sperm trajectories at different efflux rates (*j*_0_) under an exponential SAAF fields. Left: Average velocity vectors. Right; The average magnitude of the velocity vector ***v*** and its components *v*_*x*_ and *v*_*y*_ for different values of the efflux rates, *j*_0_. The visualization of velocity and mean velocity vectors is identical to that in Fig. 4C,D. a. Trajectory at *j*_0_ = 0.55. b. Trajectory at *j*_0_ = 0.6. Legends are as in Fig. 4B. B: a. Experimental ascidian sperm trajectories. Sperm swimming trajectories in low-Ca^2+^ (0.01, 0.1, and 1 mM), normal-Ca^2+^ (10 mM, control), and high-Ca^2+^ (100 mM) ASW around the tip of a capillary containing 0.1 *µ*M SAAF. The arrow indicates the direction of sperm swimming direction. b. Magnitude of radial velocity magnitude at various extracellular Ca^2+^ concentrations ([Ca^2+^]_ex_). Experimental data were obtained from Yoshida et al., 2018 [20]. Sample number *N* is indicated in bracket and each sample are obtained from three different experiments.

K. Yoshida et al. [20] investigated the effect of varying extracellular Ca^2+^ concentrations (Fig. 5B). Low extracellular Ca^2+^ concentrations correspond to low values of *j*_0_ in our model. Notably, the experimental results at 0.01 mM Ca^2+^ and case (a) in Fig. 5A exhibit similar behaviour. We analysed these experimental data to obtain the mean speed to the attractant, −*V*_*r*_. At high extracellular Ca^2+^ concentrations, the parameter *j*_0_ assumes a high values, resulting in a simulated mean trajectory that is nearly perpendicular to the SAAF gradient (case (c) in Fig. 5A). In contrast, the experimental trajectory observed at 100 mM Ca^2+^ appears disrupted, potentially due to the influence of Ca^2+^ influx under high extracellular concentrations; this influx effect was not incorporated into the present simulation. Figure 5Bb shows the distribution of the radial velocity magnitude at different extracellular Ca^2+^ concentrations. The averages peak at a certain value (10 mM), which is qualitatively consistent with our result (red curve in Fig. 5A).

## 3 Discussions

### 3.1 Role of PMCA and regulatory signals in the modelled signal transduction pathway

Our study model incorporates a Ca^2+^ pump whose activity depends on the concentration of SAAF, and Ca^2+^ channels that function independently. The pump is modelled after PMCA, which acts as a receptor for SAAF [20], whereas the channel is based on CatSper, a sperm-specific ion channel [28]. PMCA, located in the flagellar membrane, plays a critical role in maintaining Ca^2+^ homeostasis in eukaryotic cells. It contributes to both intracellular Ca^2+^ signalling and to turning behaviour during the chemotactic response to SAAF.

In this model, we introduce a signal *r* that activates the pump, and a signal *i* that inhibits it. Ascidian PMCA contains intracellular domains such as CaM and 14-3-3 [20]. As 14-3-3*ζ*, when expressed in HeLa cells, has been shown to inhibit the activity of PMCA4 ― a key isoform [29] ―, Based on this evidence, we hypothesized that SAAF-mediated activation of PMCA may also lead to the generation of an inhibitory signal, thereby establishing a negative feedback loop.

Interestingly, a similar regulatory principle underlies the well-studied mechanism of fold-change detection (FCD) in *E. coli* chemotaxis. In this system, the FCD property of the response to attractants is explained by a signal transduction model incorporating receptor methylation, in which the average kinase activity exerts inhibitory control over the methylation process. This mechanism accounts for the behavioural observations in chemotaxis experiments [12]. Although our model does not specify a detailed biochemical transduction pathway, the incorporation of a negative feedback structure suggests a potential dynamic similarity. This correspondence may provide valuable insights into the emergence of FCD-like behaviour in more abstract or generalised biological systems, and serve as a basis for future mechanistic investigations.

### 3.2 Advantages of helical trajectories for chemotaxis mechanism

From a mathematical perspective, helical or circular trajectories with nonzero curvature (and torsion) are the key advantages of the gradient-sensing algorithm, which is necessary for chemotaxis. Effective chemotaxis in three (or two) dimensions requires the detection of chemical gradients, which in turn necessitates sampling concentration values at a minimum of four non-coplanar points that form a linearly independent set (or three non-collinear points in two dimensions), which is discussed in SI 8. Points distributed along a spiral (or a circle in two dimensions) generally satisfy this condition. By contrast, a straight trajectory represents the minimal path between two points and may appear to be a more optimal geometric shape than a spiral (or a circle). However, any four points along a straight trajectory are linearly dependent, which renders gradient detection impossible.

Therefore, helical (or spiral) trajectories provide informational adequacy, making them particularly advantageous for gradient sensing and chemotactic navigation.

The present model describes a chemotaxis algorithm employed by ascidian sperms, which can be interpreted as a gradient detection algorithm. Compared to the method of steepest descent method, our method exhibits greater robustness to noise because it integrates chemical concentration information along the sperm trajectory to infer the gradient. A rough estimate suggests that the diameter of a typical circular trajectory approximates the length scale of the gradient, which is the case demonstrated in Fig. 2E. This type of algorithm is likely to be advantageous in natural environments where noise is prevalent.

### 3.3 FCD in ascidian sperm trajectory

To the best of our knowledge, this is the first report demonstrating FCD property in ascidian sperm. The trajectories under the different level of SAAF field but proportional to one another showed similar shape and radial velocity (Fig.2C, SI 7). Although FCD has been suggested to be within a certain range of SAAF concentrations, the precise boundaries of this effective range have not yet been fully determined. Notably, SAAF may function not only as a chemoattractant but also as an activator of sperm motility. From a biological perspective, FCD is advantageous in natural environments where eggs are subject to water currents and the distribution of attractants is perturbed by other sperm, requiring sperm to effectively locate their targets. Our results provide insights into the mechanisms underlying the navigation of sperms towards eggs.

In our model, the averaged trajectory was constant under the expoenetial distribution of the attractant, and this distribution was used to analyse the chemotaxis ability (Fig.4). In sea urchin sperm, the spatial decay rate of attractants is the key to inducing chemotaxis [30]. Our analysis suggests the mathematical properties of the signalling inside the cell by focusing on the trajectory shape and FCD properties.

### 3.4 Relative gradient shapes the canonical chemotactic trajectory

We characterised sperm trajectories under the attractant distribution *s*(***x***) using the relative decay rate *α*, and found that chemotactic efficiency defined as the maximum velocity component aligned with the concentration gradient achieves its peak at an optimal value of *α* (see Fig. 4D).

A simple linear gradient, in which the gradient of *s*(*x*) is constant throughout space, does not lead to efficient trajectories, as the relative decay rate *α* varies spatially (see Fig. 4A).

Notably, in experiments using rectangular agar soaked with SAAF, the resulting attractant distribution —described by a diffusion equation with Dirichlet boundary conditions at the agar-solution interface— is concave. Although it is not strictly exponential, the spatial variance of *α* is smaller than in the linear gradient case. This probably explains the linearity of the period-averaged trajectories, observed in the experiments in Fig. 4E.

The model reproduces key features of sperm trajectories for different attractant profiles, including radial distribution and one-dimensional distributions. Furthermore, the emergence of linear period-averaged trajectories under exponential distributions is a direct consequence of the FCD properties (see SI 6).

## Acknowledgments

This work was supported by MEXT KAKENHI Grant-in-Aid for Transformative Research Areas (A), “Ethological dynamics in diorama environments”, (21H05303, 21H05311, 21H05304) and for Grant-in-Aid for Scientific Research (B) (25291069). *Ciona* adults were provided by Satou’s lab, (Kyoto University) and Yoshida’s lab (University of Tokyo) under the support of the National BioResource Project, MEXT, Japan, and the staff members of Onagawa Field Center, Graduate School of Agricultural Science, Tohoku University.

## Author Contributions

M.I. and K.S. designed the study and wrote the manuscript. M.I. constructed mathematical model, performed theoretical analyses. K.S. and K.I. performed experiments and related analyses. M.Y. gave experimental data. M.I, K.S., K.I. and T.N. managed funding acquisition. T.N. supervised the project. All authors contributed to discussions and review of the manuscript.

## 4 Methods

In this study, we examined sperm motion in a two-dimensional plane, reflecting the experimental observation that ascidian sperm predominantly move near surfaces. The experimental details are described in SI 1. The proposed mathematical model consists of two main components: (I) signal transduction from SAAF to the intracellular Ca^2+^ concentration, [Ca^2+^]_*i*_, and (II) sperm motility governed by curvature, which is determined by the instantaneous value of [Ca^2+^]_*i*_. The details are described in SI 2.

In Part (I), the receptor response is expressed as follows. Let *s* denote the SAAF concentration at the representative position of the sperm. The receptor generates a signal *r*, an internal variable as an inhibitor *i*, governed by the following equations:

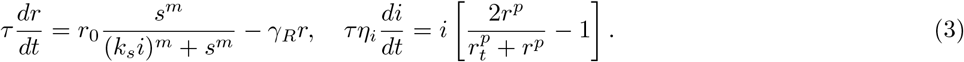

These equations collectively describe a negative feedback system. Furthermore, this system shows FCD properties by assuming *s* as the input, *r* as the output, and *i* as the internal variable [10].

For the Ca^2+^ pump, [Ca^2+^]_*i*_ (denoted as *c*) are expressed as 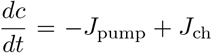, where *J*_pump_ represents the flux of Ca^2+^ pumped out of the cell, and *J*_ch_ represents the flux of Ca^2+^ entering through the channels (Fig. 1A). The dynamics of *c* are expressed as:

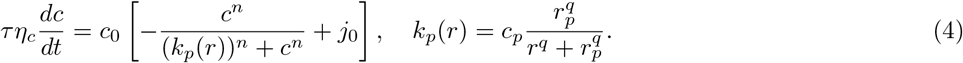

We assumed that the Ca^2+^ pump was represented by the Hill equation [31, 32, 33, 34, 35].

In part (II), the sperm is modelled as a point located at *x* = (*x, y*), moving with a velocity *v* = (*v*_*x*_, *v*_*y*_) at a constant speed |*v*| = *v*_0_. The curvature of the sperm’s trajectory, *κ*, is assumed a function of *c*: 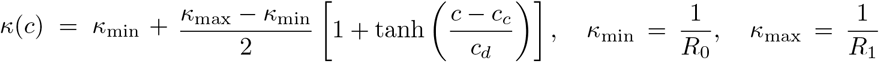. Here *c*_*c*_ and *c*_*d*_ are constants, while *R*_0_ and *R*_1_ represents the asymptotic radius of curvature when *c* ≪ *c*_*c*_ and *c* ≫ *c*_*c*_, respectively. For simplicity, the model omits the detailed response process observed in ascidian sperms, in which they swim with a large curvature around regions of locally minimal SAAF concentration, transition to nearly straight swimming, and subsequently recover their curvature [21].

Once the curvature *κ* is determined, the sperm’s motion is described by Frenet equations:

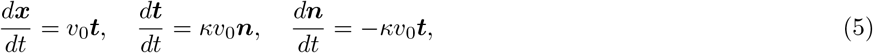

where, *t* and *n* are the tangential and normal unit vectors of the trajectory, respectively. The period *T* when *c* ≪ *c*_*c*_ can be related to the radius *R*_0_ as 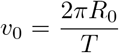. In the three-dimensional model (SI 3), the sperm motion is described by Frenet-Seret equation with a constant torsion of 0.01.

