## Supplementary Information for "Spiral-Sensing and Fold-Change Detection Direct Ascidian Sperm to the Egg"

#### 1 Experimental details

The ascidian *Ciona intestinalis* (type A; also called *C. robusta*) was collected from Onagawa Bay near the Onagawa Field Research Center, Tohoku University or obtained from the National BioResource Project for *Ciona* (`https://marinebio.nbrp.jp`). The animals were maintained in aquaria under constant light condition for gemete accumulation without spontaneous spawning. Semen samples were collected by dissecting the sperm ducts and were stored on ice until use. Artificial seawater (ASW) was composed of 462.01 mM NaCl, 9.39 mM KCl, 10.81 mM $CaCl_2$, 48.27 mM $MgCl_2$ and 10 mM HEPES-NaOH (pH 8.0). SAAF was synthesised as previously described [1, 2] . High and low $Ca^{2+}$-ASW were prepared as the ASW but with 0.01, 0.1, 1, 50 or 100 mM $CaCl_2$ and 5 mM EGTA. Sperm chemotaxis was examined as previously described [3] . Briefly, semen was suspended in 2000 volumes of ASW with SAAF. After incubation for 1 min, the sperm suspension was placed in an observation chamber made of a 0.5 mm silicone rubber sheet (6-611-01, ASONE, Osaka, Japan), and sperm movement around the micropipette tip containing SAAF in ASW with 1% agar was observed under a phase contrast microscope (BX51, Olympus, Tokyo, Japan) with a 10x objective (UPlan FL N, Olympus) and recorded using a high-speed CCD camera (HAS220 or HASU2, DITECT, Tokyo, Japan) at 100 or 200 fps. For analysis in a 1D linearly distributed SAAF field, half of the chamber was filled with 1 µM SAAF in ASW with 1% agar and the sperm suspension was placed to the other side. The observations and recordings were performed within 5 minutes of sperm dilution. The position of the sperm head was analyzed using Bohboh software (BohbohSoft, Tokyo, Japan). All experiments were repeated at least three times with three different animals. Statistical significance was calculated using Dunnett's test; $p < 0.05$ was considered significant. Data were analysed by using the R script.

#### 2 Model details

The proposed mathematical model consists of (I) signal transduction from SAAF to intracellular $Ca^{2+}$ concentration. (II) Sperm motility is governed by curvature, which is determined by the instantaneous value of $[Ca^{2+}]_i$.

In part (I), the functions of the receptor response and $Ca^{2+}$ pumping by PMCA are segregated. The $Ca^{2+}$ pump was modelled as a single entity that aggregates the calcium transport roles of PMCA (Fig. 1A). In Eq. (3), the generation of $r$ is described by a Hill equation with respect to $s$, while its decay is modeled as a linear process with a timescale of $\tau/\gamma_R$. As the Hill equation is an increasing function of $s$, yielding 1/2 when $k_s i = s$, a higher $i$ suppresses the level of $r$. The generation of the inhibitor $i$ is also governed by a Hill equation with respect to $r$, coupled with linear decay with a timescale of $\tau\eta_i$. Specifically, $i$ increases (decreases) when $r > r_0$ ($r < r_0$), where $r_0$ is a constant.

For the $Ca^{2+}$ pump, the equation of the type $\dfrac{dc}{dt} = -J_{\rm pump} + J_{\rm ch}$ is assumed. Here $J_{\rm pump}$ represents the flux of $Ca^{2+}$ pumped out of the cell, and $J_{\rm ch}$ represents the flux of $Ca^{2+}$ entering through channels (Fig. 1A).

The pump flux, $J_{\rm pump}$, is given by $J_{\rm pump} = V_p \frac{c^n}{(k_p(r))^n + c^n}$ [4, 5, 6, 7, 8], where $k_p(r)$ represents the activation threshold of the pump, and a higher $r$ decreases this threshold, thereby enhancing $J_{\rm pump}$. The threshold $k_p(r)$ follows: $k_p(r) = c_0 \frac{r_e^m}{r^m + r_e^m}$. The channel flux, $J_{\rm ch}$, is modeled as diffusion $J_{ch} = K_2(c_e - c)$, where $c_e$ is the extracellular $Ca^{2+}$ concentration where $K_2$ is a constant. Given $c_e \gg c$, $J_{\rm ch} \approx J_0$ (constant).

By combining these elements, we obtained Eq. (4), in which the constants are replaced with simplified expression.

In summary, an increase in $s$ activates $r$, which subsequently increases $i$. Elevated $r$ enhances the activity of the $Ca^{2+}$ pump, reducing $c$. Simultaneously, the rise in $i$ suppresses $r$, leading to decreased pump activity and allowing $Ca^{2+}$ influx to dominate. An oscillatory input of $s$ induces a phase-delayed oscillation in $c$, whereby enabling an anti-phase relationship.

### 3 Simulation conditions

The model parameters are listed in Table 1. The non-dimensional value for parameters are provided by the length scale $L = 50$µm, the time scale 0.5s and the concentration scale 1nM. The values in the main text and SI are written in the dimensional form.

| Equations | Constant | Variable | Unit | Non-dimensional value |
|---|---|---|---|---|
| Receptor (Eq.(3)) | Maximum level of $r$ | $r_0$ | nM | 1 |
| | Time scale of $r$ production | $\tau$ | s | 0.08 |
| | Hill coefficient | $m$ | - | 2 |
| | Decay rate constant of $r$ | $\gamma_R$ | 1 | 0.5 |
| | Inhibition constant | $k_s$ | 1 | 5.0 |
| Inhibitor (Eq.(3)) | Relative time scale of $i$ production to $\tau$ | $\eta_i$ | 1 | 3.3 |
| | Hill coefficient | $p$ | - | 2 |
| | Threshold concentration of $r$ for $i$ production | $r_t$ | nM | 0.6 |
| $Ca^{2+}$ pump (Eq.(4)) | Maximum level of $c$ | $c_0$ | nM | 1 |
| | Relative time scale of the pump to $\tau$ | $\eta_c$ | 1 | 3.3 |
| | Hill coefficient of the pump dynamics | $n$ | - | 2 |
| | Maximum value of $k_p$ | $c_p$ | nM | 8.0 |
| | Hill constant of the threshold | $q$ | - | 2 |
| | Half-saturation concentration of $r$ for $k_p$ | $r_p$ | nM | 0.11 |
| | Relative flux into $Ca^{2+}$ channel | $j_0$ | 1 | 0.6 |
| Sperm dynamics (Eq.(5)) | Period | $T$ | s | 3.6 |
| | Maximum radius | $R_0$ | µm | 1.2 |
| | Minimum radius | $R_1$ | µm | 0.4 |
| | Threshold of the variation of $Ca^{2+}$-curvature | $c_c$ | nM | 0.35 |
| | Width of the variation of $Ca^{2+}$-curvature | $c_d$ | nM | 0.02 |
| Sperm dynamics in 3D | Torsion | $\chi$ | 1/µm | 5 |

Table 1: List of model parameters.

The simulation code was developed using Python 3.12.3 with the `SciPy` and `Numpy` libraries. Ordinary differential equations were integrated using the `scipy.integrate.odeint` function, with the error tolerance parameters set to `rtol=atol=1e-10`.

For the signal transduction process described by Eqs. (3) and (4) under the given function of $s(t)$ defined in Sec. 2.1, Figs. 1B and 1C were generated under the following conditions: $s_{\rm max} - s_{\rm min} = 5.0$, $s_{\rm min} = 1.0$, and $\omega = \frac{2\pi}{0.5}$. The period of $s(t)$ was set to 0.5, which is a typical value observed in ascidian sperm experiments, and $s_{\rm max} - s_{\rm min} = 5.0$ was chosen to clearly illustrate the phase delay in the signal transduction system.

The initial conditions for Eqs. (3) and (4) were set as $i = r = c = 0.5$. The trajectories of $s$ and $c$ (Fig. 1C) were drawn using data from the time interval $t > T_{\rm end}/2$, where $T_{\rm end} = 3.0$.

For the chemotactic behavior described by Eqs. (3), (4), and (5) under the given field $s(\boldsymbol{x})$ defined by Eq. (1), the initial conditions for the particle position and orientation are specified as:

$$\boldsymbol{x}(0) = (x_0, y_0), \quad \boldsymbol{t}(0) = (\cos\theta, \sin\theta), \quad \boldsymbol{n}(0) = (-\sin\theta, \cos\theta), \quad (x_0, y_0, \theta) = (400, -400, 0).$$

The curvature $\kappa = 1/R_0$ was maintained during the time interval $t \leq 3T$ to allow stabilisation of the signal transduction reactions. The full dynamics were computed for $t > 3T$.

When noise was introduced into the field $s(\boldsymbol{x})$, the field was defined as:

$$s(\boldsymbol{x}) = s_{\max}(1 + A\eta(\sigma, \ell; \boldsymbol{x}))f(\boldsymbol{x}, \boldsymbol{x}_0; \epsilon),$$

where $f(\boldsymbol{x}, \boldsymbol{x}_0; \epsilon)$ is defined in Eq.(1), $\alpha$ is the relative amplitude. The noise $\eta(\sigma, \ell; \boldsymbol{x}))$ follows the Gaussian process of zero mean,

$$\eta(\sigma, \ell; \boldsymbol{x})) \sim GP(0, k(\sigma, \ell; \boldsymbol{x}, \boldsymbol{x}')),$$

where $k(\sigma, \ell; \boldsymbol{x}, \boldsymbol{x}')$ is the radial basis function, i.e.,

$$k(\sigma, \ell; \boldsymbol{x}, \boldsymbol{x}') = \sigma^2 \exp\left(-\frac{|\boldsymbol{x} - \boldsymbol{x}'|^2}{2\ell}\right).$$

In the simulation, we set $A = 0.1$, $\sigma = 1$ and controlled the correlation length $\ell$.

In the three-dimensional model, Eq.(5) is replaced by the Frenet–Serret equations,

$$\frac{d\boldsymbol{x}}{dt} = v_0\boldsymbol{t}, \quad \frac{d\boldsymbol{t}}{dt} = \kappa v_0\boldsymbol{n}, \quad \frac{d\boldsymbol{n}}{dt} = v_0(-\kappa\boldsymbol{t} + \chi\boldsymbol{b}), \quad \frac{d\boldsymbol{b}}{dt} = -\chi v_0\boldsymbol{n}. \tag{SI 1}$$

where $\chi$ and $\boldsymbol{b}$ denote the torsion and the binormal unit vector of the trajectory, respectively. The simulation was carried out with the initial condition $\boldsymbol{x}_0 = (350, 350, -350)$, and the initial frame $\boldsymbol{t}(0)$, $\boldsymbol{n}(0)$, and $\boldsymbol{b}(0)$ was determined based on the axis of a helix with constant curvature $1/R_0$ and torsion $\chi$ (see SI 6).

### 4 An extension of fold-change detection

We discuss the fold-change detection properties and an extension to the case with plural output variables. First, we consider the dynamical system

$$\frac{dx}{dt} = f(x, y; z), \quad \frac{dy}{dt} = g(x, y; z), \tag{SI 2}$$

where $z$ represents the input, $x$ the internal variable and $y$ the output.

The outputs $y_1(t), y_2(t)$ given by the two inputs $z = z_1(t), z_2(t) = pz_1(t)$ ($p > 0$ is a constant), are said to be fold-change detection (FCD) when $y_1(t) = y_2(t)$. [9]

A condition to hold FCD property for $f$ and $g$ are:

$$f(px, y; pz) = pf(x, y; z), \quad g(px, y; pz) = g(x, y; z), \tag{SI 3}$$

[10].

Here we consider the case with multiple output variables. Let us consider the following dynamical systems with $(n+1)$ dimensions,

$$\frac{dx_j}{dt} = f_j(x_0, \cdots, x_n; z), \quad (j = 0, 1, \cdots, n), \tag{SI 4}$$

where $z$ denotes the input, $x_0$ denotes the internal variable and $x_k$ $(k = 1, \cdots, n)$ denotes the output of the $n$ variable.

**Theorem 1**:

If there exists a function $\phi(p, x_0)$ such that, for any $p > 0$, the following relationships

$$f_0(\phi(p, x_0), x_1, \cdots, x_n; pz) = \frac{\partial \phi}{\partial x_0} f(x_0, x_1, \cdots, x_n; z), \quad f_j(\phi(p, x_0), x_1, \cdots, x_n; pz) = f_j(x_0, x_1, \cdots, x_n; z), (j = 1, \cdots, n) \tag{SI 5}$$

hold, then the dynamical system (SI 4) satisfies FCD property, that is, the outputs $\{x_j^{(1)}(t) \mid j = 1, \cdots, n\}$ and $\{x_j^{(2)}(t) \mid j = 1, \cdots, n\}$ given by the two inputs $z = z^{(1)}(t)$ and $z = z^{(2)}(t) = pz^{(1)}(t)$ are the same: $x_j^{(1)}(t) = x_j^{(2)}(t)$ $(j = 1, \cdots, n)$. In particular, the case that $\phi(p, x) = px$ and $n = 1$ is equivalent with the system determined by Eqs. (SI 2) and (SI 3).

**Proof**:

Let $f_j$ satisfy the condition (SI 5). Assume that two inputs $z^{(1)}(t)$ and $z^{(2)}(t)$ satisfies $z^{(2)}(t) = pz^{(1)}(t)$, that the corresponding internal variables are $x_0^{(1)}(t)$ and $x_0^{(2)}(t)$, and that corresponding output variables are $\{x_j^{(1)}(t)\}$ and $\{x_j^{(2)}(t)\}$ $(j = 1, \cdots, n)$. Then the following two equations hold:

$$\frac{dx_j^{(1)}}{dt} = f_j(x_0^{(1)}, \cdots, x_n^{(1)}; z^{(1)}), (j = 0, \cdots, n), \tag{SI 6}$$

$$\frac{dx_j^{(2)}}{dt} = f_j(x_0^{(2)}, \cdots, x_n^{(2)}; z^{(2)}), (j = 0, \cdots, n). \tag{SI 7}$$

We show that $\{x_0^{(2)}, x_1^{(2)}, \cdots, x_n^{(2)}\} = \{\phi(p, x_0^{(1)}), x_1^{(1)}, \cdots, x_n^{(1)}\}$ satisfies Eq.(SI 7).

Using the relationship (SI 5), we can show the following:

$$\begin{aligned}
\frac{dx_0^{(2)}}{dt} - f_0(x_0^{(2)}, \cdots, x_n^{(2)}; z^{(2)}) &= \frac{\partial \phi}{\partial x_0}(p, x_0^{(1)})\frac{dx_0^{(1)}}{dt} - f(\phi(p, x_0^{(1)}), x_1^{(1)}, \cdots, x_n^{(1)}; pz^{(1)}) \\
&= \frac{\partial \phi}{\partial x_0}(p, x_0^{(1)}) \left( \frac{dx_0^{(1)}}{dt} - f(x_0^{(1)}, x_1^{(1)}, \cdots, x_n^{(1)}; z^{(1)}) \right) = 0. \\
\frac{dx_k^{(2)}}{dt} - f_k(x_0^{(2)}, \cdots, x_n^{(2)}; z^{(2)}) &= \frac{dx_k^{(1)}}{dt} - f_k(\phi(p, x^{(1)}), x_1^{(1)}, \cdots, x_n^{(1)}; pz^{(1)}) \\
&= \frac{dx_k^{(1)}}{dt} - f_k(x_0^{(1)}, x_1^{(1)}, \cdots, x_n^{(1)}; z^{(1)}) = 0. \quad (k = 1, 2, \cdots, n)
\end{aligned}$$

□

In the signal transaction system, $n = 4$ and assuming $z = s(t), x_0 = i, x_1 = r, x_2 = c$, Eqs.(1) and (2) satisfy the following condition (SI 4).

In this chemotactic model, $n = 5, z = s(\boldsymbol{x}(t)), x_0 = i, x_1 = r, x_2 = c, x_3 = x, x_4 = y, x_5 = \theta_t$, where $\boldsymbol{t} = (\cos\theta_t, \sin\theta_t)$ and $\boldsymbol{n} = (-\sin\theta_t, \cos\theta_t)$. and $\phi(p, x) = px$, the conditions of the theorem are satisfied, so the FCD is satisfied. In other words, as $s(\boldsymbol{x}) \to \alpha s(\boldsymbol{x})$, the variables except $i$, in particular the internal calcium concentration $c$ and the orbit $\boldsymbol{x}(t)$, remain unchanged.

### 5 Robustness of the signal transduction model

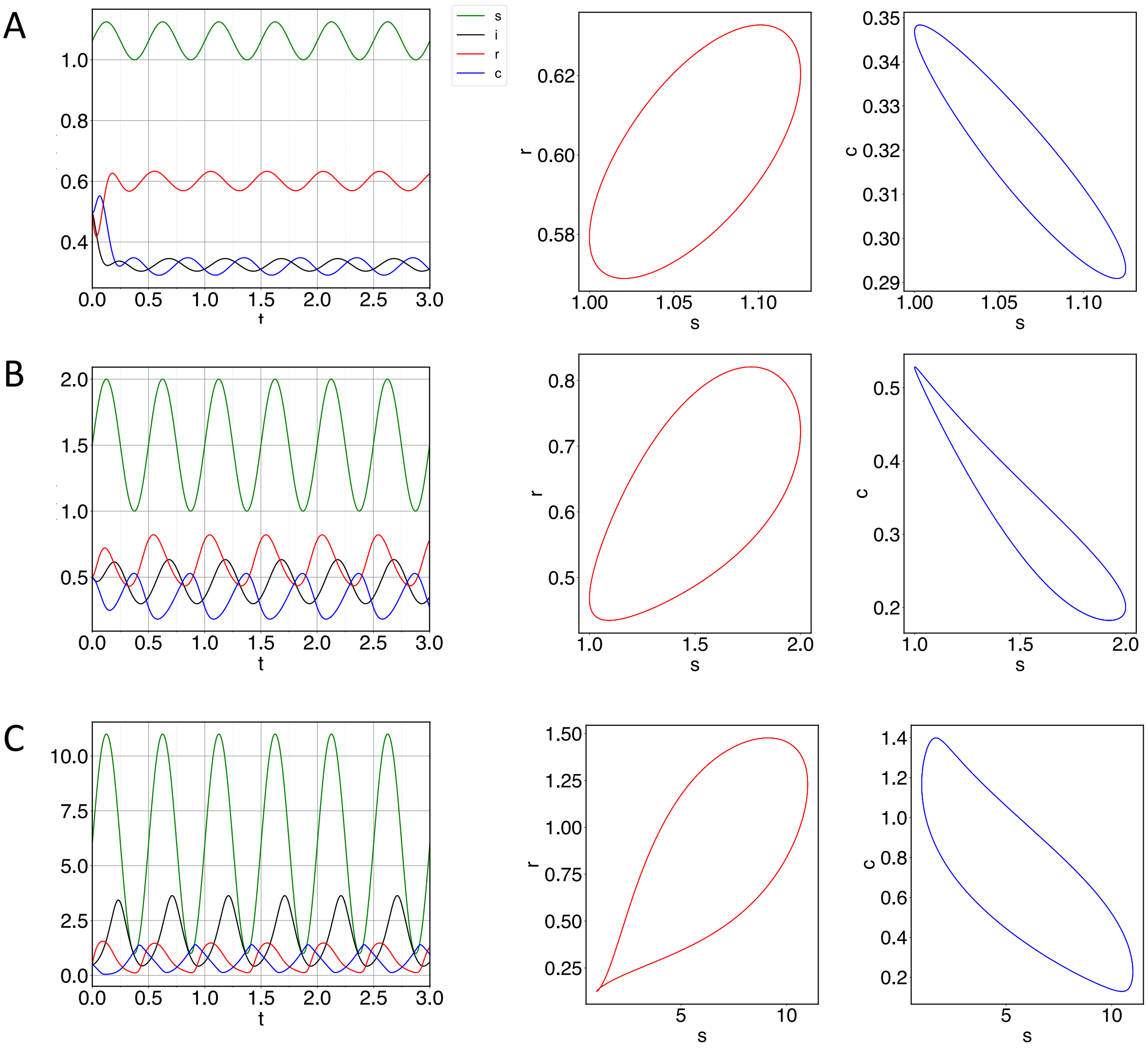


Figure 1: A. Signal transduction system with $s_{\max} - s_{\min} = 0.125$. Left: Time series of $s$, $i$, $r$, and $c$. Middle: Trajectory in the $(s, r)$ plane. Right: Trajectory in the $(s, c)$ plane. B. Same as (A), but with $s_{\max} - s_{\min} = 1.0$. C. Same as (A), but with $s_{\max} - s_{\min} = 10.0$.

Figure 1 illustrates the responses to the oscillatory signal $s(t) = s_{\mathrm{osc}}(t)$ in the signal transduction system (Eqs. (3) and (4) in the main text) for different values of $s_{\max} - s_{\min}$. Here, the oscillation period was set to 0.5, which is comparable to the rotational period observed in the chemotaxis simulation (Fig. 2). In the chemotaxis simulation, the period was estimated to be 0.63 s in the range $5 \le t \le 10$ and 0.45 s in the range $10 \le t \le 15$.

Due to the FCD property of the signal transduction system (see SI 4), the output signal $c$ remains invariant when $(s_{\min}, s_{\max})$ is replaced by $(\alpha s_{\min}, \alpha s_{\max})$, where $\alpha$ is a constant. Therefore, without loss of generality, we assume $s_{\min} = 1$.

As demonstrated in the three cases of $s_{\max} - s_{\min} = 0.125, 1.0$, and 10.0, the in-phase relationship between $s$ and $r$ and the anti-phase relationship between $s$ and $c$ are maintained across a wide range of $s_{\max} - s_{\min}$ values (spanning nearly by two orders of magnitude). However, the amplitude of $c$ increases as $s_{\max} - s_{\min}$ increases.

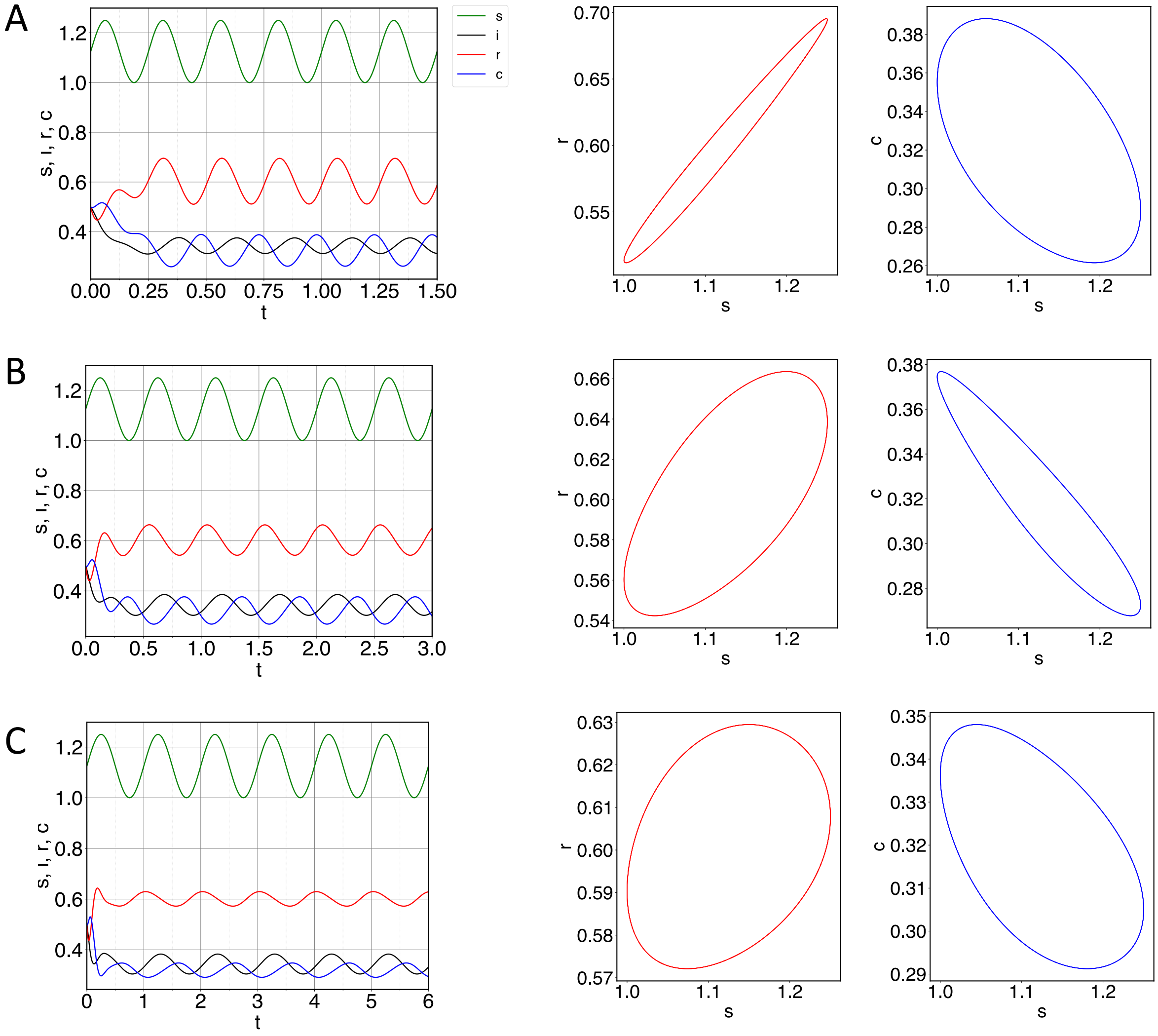


Figure 2: A Signal transduction system with $s_{\min} = 1.0$, $s_{\max} - s_{\min} = 0.25$, and $\omega = 2\pi/0.25$. Left: Time series of $s$, $i$, $r$, and $c$. Middle: Trajectory in the $(s, r)$ plane. Right: Trajectory in the $(s, c)$ plane. B Same as A, but with $\omega = 2\pi/0.5$. C Same as A, but with $\omega = 2\pi/1.0$.

Figure 2 illustrates the responses of the signal transduction system to the input signals $s_{\rm osc}(t)$ for varying periods. The input signal parameters were set to $s_{\min} = 1.0$ and $s_{\max} - s_{\min} = 0.25$. The examined oscillation periods were 0.25, 0.50, and 1.0. Within this periods range, both the in-phase relationship between $s$ and $r$ and the anti-phase relationship between $s$ and $c$ are maintained. Given that the typical period of the input signal in the experiments, $T_0$, was approximately 0.5, these results demonstrate the robustness of the antiphase relationship between $s$ and $c$ for input signals with periods ranging from half ($T_0/2$) to double ($2T_0$) the typical period. However, that the antiphase relationship is lost for periods outside this range.

#### 6 Robustness of the sperm trajectory

In the sperm model, the success rate of navigation towards the egg (defined as reaching the high-concentration region, $|\boldsymbol{x} - \boldsymbol{x}_0| < \epsilon$ at $t = T_{\rm end} = 20$) was evaluated ($N = 20$). The observed success rates were 19/20, 20/20, 17/20, and 15/20 for $(l, \alpha) = (10, 0.05)$, $(25, 0.05)$, $(10, 0.1)$, and $(25, 0.1)$, respectively, indicating a consistently high level of successful navigation despite the presence of various types of noise.

In the three-dimensional model, the success rate, defined as $|\boldsymbol{x} - \boldsymbol{x}_0| < 1.5\epsilon$ at $t = T_{\rm end} = 20$, was similarly evaluated ($N = 100$). A trajectory with constant curvature and torsion describes a helix that possesses a well-defined axis $\boldsymbol{a}$ along which the trajectory tends to extend on average. When the initial direction of $\boldsymbol{a}$ was randomly selected, the overall success rate was 67%, despite the fact that it included directions oriented away from the source.

### 7 Orbit invariance under exponential SAAF distribution

We considered the condition of $s(\boldsymbol{x})$ for the chemotaxis system, in which the sperm velocity per period was constant. For this purpose, let $\boldsymbol{Y} = (r, c, \boldsymbol{t}, \boldsymbol{n})^t$, the output variables of the chemotaxis system excluding the position $\boldsymbol{x}$.

Assume that a trajectory satisfies $\boldsymbol{Y}(t+T) = \boldsymbol{Y}(t)$ and $\boldsymbol{x}(t+T) - \boldsymbol{x}(t) = \boldsymbol{d}$ for any $t$, where $\boldsymbol{d}$ is a constant vector. The trajectory is called "constant drift trajectory", $\boldsymbol{d}$ is called "the drift", and $T$ is called "the period".

**Theorem 2**:
Suppose that $\boldsymbol{Y}(0) = \boldsymbol{Y}(T)$ for $T$ and $s(\boldsymbol{x})$ is represented as

$$s(\boldsymbol{x}) = A \exp(-\alpha \hat{\boldsymbol{u}} \cdot \boldsymbol{x}), \tag{SI 8}$$

where $A$ is a constant and $\hat{\boldsymbol{u}}$ is a constant unit vector. Subsequently, the orbit exhibits a constant drift trajectory.

**Proof**:
For any position vector $\boldsymbol{x}$, any constant vector $\boldsymbol{q}$ yields

$$s(\boldsymbol{x}+\boldsymbol{q}) = A \exp\left(-\alpha \hat{\boldsymbol{u}} \cdot (\boldsymbol{x}+\boldsymbol{q})\right) = \exp\left(-\alpha \hat{\boldsymbol{u}} \cdot \boldsymbol{q}\right) s(\boldsymbol{x}) = \beta(\boldsymbol{q}) s(\boldsymbol{x}) \quad (\beta(\boldsymbol{q}) = \exp\left(-\alpha \hat{\boldsymbol{u}} \cdot \boldsymbol{q}\right) \text{is a constant}). \tag{SI 9}$$

Define $\boldsymbol{d}_0 = \boldsymbol{x}(T) - \boldsymbol{x}(0)$ and consider a trajectory defined by

$$\boldsymbol{x}(t) = \boldsymbol{x}(t-T) + \boldsymbol{d}_0, \boldsymbol{Y}(t) = \boldsymbol{Y}(t-T) \quad (T \leq t \leq 2T). \tag{SI 10}$$

Then, the trajectory (SI 10) is the solution of the model. This is expressed as follows. 1) The input signal along the trajectory is determined using Eq. (SI 10), $s(\boldsymbol{x}(t))$ $(T \leq t \leq 2T)$ satisfies $s(\boldsymbol{x}(t)) = s(\boldsymbol{x}(t-T) + \boldsymbol{d}_0) = \beta(\boldsymbol{d}_0) s(\boldsymbol{x}(t-T))$, according to Eq. (SI 9); thus, the time series of $s$ is folded by a constant $\beta(\boldsymbol{d}_0)$. 2) The FCD property indicates that the trajectory (SI 10) is a solution for the chemotaxis system. 3) The uniqueness of an ordinary differential equation guarantees that its solution is unique. Similar discussion leads that the trajectory of the sperm trajectory under the field $s(\boldsymbol{x})$ satisfies $\boldsymbol{x}(t+nT) = \boldsymbol{x}(t) + n\boldsymbol{d}_0$ $(0 \leq t \leq T,\ n \in \mathbb{Z})$, which means the trajectory is the constant drift trajectory. □

### 8 FCD experiment

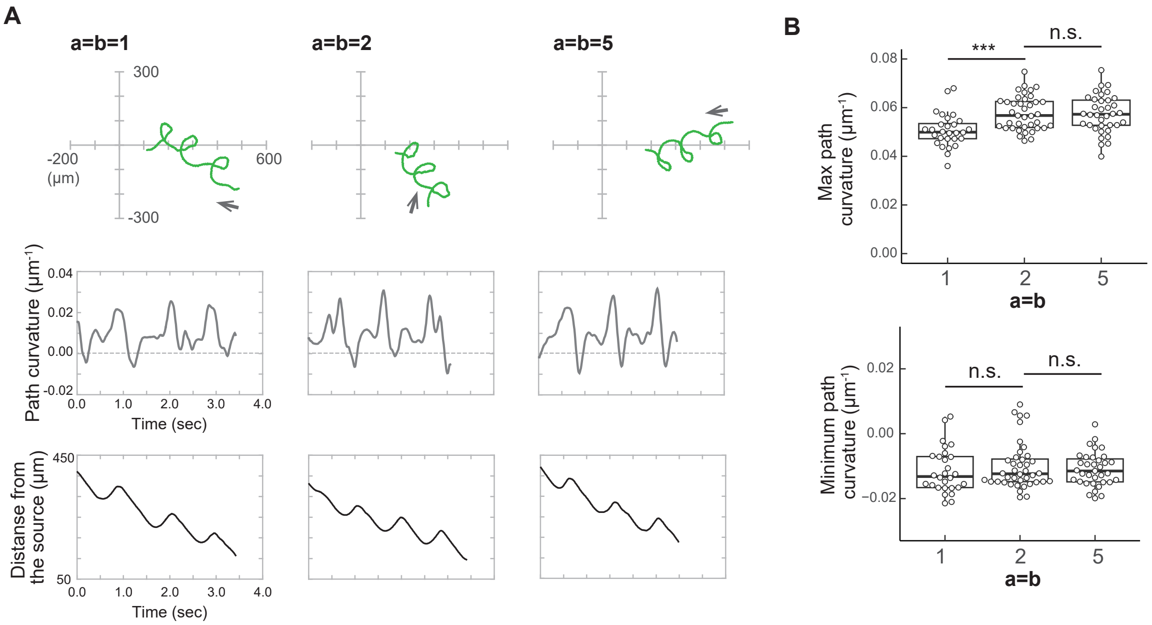


Figure 3: A: Typical trajectories under the condition $(a, b) = (1, 1), (2, 2)$ and $(5, 5)$. Top: Sperm swimming trajectories in physical space. Arrows indicate the direction of swimming. Middle: Time series of the curvatures. Bottom: Time sereis of the distances from the attractant. B: The maximum and minimum curvatures of between turns when the sperm approaches the source. Sample numbers N = 28-39 from three different experiments. Distribution of values is plotted in a box plot. *** Significant at $p < 0.001$ (Dunnet test), as compared with the value of (a, b) = (2, 2).

The definition of $V_r$ in the main text (Fig. 3Ab) is as follows: The characteristic time of $k$th turn, $t_k$, was defined as the time when the distance from the origin reached the $k$th local maximum. The position of the sperm head at $t = t_k$ was defined as $\boldsymbol{r}_k = \boldsymbol{r}(t_k)$. Then, the displacement and the period between $k$th turn and $(k+1)$th turn are defined by $\boldsymbol{r}_{k+1} - \boldsymbol{r}_k$ and $t_{k+1} - t_k$, respectively. Mean velocity between $k$th turn and the $(k+1)$th turn, $\boldsymbol{v}^{(k)}$, is defined by $\boldsymbol{v}^{(k)} = (\boldsymbol{r}_{k+1} - \boldsymbol{r}_k)/(t_{k+1} - t_k)$, and its radial component $v_r^{(k)}$ is defined by $v_r^{(k)} = \boldsymbol{v}_k^{(k)} \cdot \boldsymbol{e}_r$, where $\boldsymbol{e}_r$ is the unit vector in radial direction. The sign of $v_r^{(k)}$ is negative if the head moves closer to the origin in this period, as the typical trajectories approaching the attractants. Figure 3A shows typical trajectories (upper figures), time series of the path curvature (middle graphs), and time series of the distance from the source (lower graphs), for three different conditions; $(a, b) = (1, 1), (2, 2)$ and $(5, 5)$ (see the captions of Figure 3).

Figure 3B shows the local maximum values of the curvature and the local minimum values of the curvature, which characterise the trajectory shape. In all cases, the values did not show significant statistical differences.

### 9 Gradient determination and points on the trajectory

We consider the condition to determine the gradient of the concentration field $c(\boldsymbol{x})$ by the values of $c$ at different $N$ points that is close to each other. The concentration field near a point $\boldsymbol{x}$, $\boldsymbol{x} + \Delta\boldsymbol{x}$, is represented by

$$c(\boldsymbol{x} + \Delta\boldsymbol{x}) = c(\boldsymbol{x}) + \Delta\boldsymbol{x} \cdot \nabla c(\boldsymbol{x}) + O(|\Delta\boldsymbol{x}|^2). \qquad \text{(SI 11)}$$

In $n-$dimensional space ($n = 2, 3$), $\nabla c$ has $n$ components; thus, $n$ independent equations are needed to determine $\nabla c$. We omit $O(|\Delta\boldsymbol{x}|^2)$ to simplify Eq.(SI 11) to $\Delta\boldsymbol{x} \cdot \nabla c(\boldsymbol{x}) = c(\boldsymbol{x} + \Delta\boldsymbol{x}) - c(\boldsymbol{x})$, which gives a linear relationship among the components of $\nabla c(\boldsymbol{x})$ from $c(\boldsymbol{x}), c(\boldsymbol{x} + \Delta\boldsymbol{x})$. For instance, in the two-dimensional case, we require equations for two different $\Delta\boldsymbol{x}_1$ and $\Delta\boldsymbol{x}_2$ that are linearly independent. Under the condition we obtain the following simultaneous linear equations for $\nabla c = (\frac{\partial c}{\partial x}, \frac{\partial c}{\partial y})$:

$$A \begin{pmatrix} \frac{\partial c}{\partial x} \\ \frac{\partial c}{\partial y} \end{pmatrix} = \begin{pmatrix} c(\boldsymbol{x} + \Delta\boldsymbol{x}_1) - c(\boldsymbol{x}) \\ c(\boldsymbol{x} + \Delta\boldsymbol{x}_2) - c(\boldsymbol{x}) \end{pmatrix}, \quad A = \begin{pmatrix} \Delta x_1 & \Delta y_1 \\ \Delta x_2 & \Delta y_2 \end{pmatrix},$$

where $\Delta\boldsymbol{x}_i = (\Delta x_i, \Delta y_i)$ $(i = 1, 2)$. The necessary and sufficient condition for the equations to have non-trivial solution is that the matrix $A$ has the inverse matrix, which is equivalent to the condition that $\Delta\boldsymbol{x}_1, \Delta\boldsymbol{x}_2$ are linearly independent. The number of the point for this condition is three, $\boldsymbol{x}, \boldsymbol{x} + \Delta\boldsymbol{x}_1, \boldsymbol{x} + \Delta\boldsymbol{x}_2$, which makes a triangle. Therefore, we selected these points on circle. When $n = 3$, similar discussion leads to the condition that four points $\boldsymbol{x}, \boldsymbol{x}+\Delta\boldsymbol{x}_1, \boldsymbol{x}+\Delta\boldsymbol{x}_2, \boldsymbol{x}+\Delta\boldsymbol{x}_3$, which makes a tetrahedron, and we can choose these points on a helix ($\Delta\boldsymbol{x}_1, \Delta\boldsymbol{x}_2, \Delta\boldsymbol{x}_3$ are linearly independent). Note that three (or four) points lying on a straight line, which represent the minimal path between two points, do not satisfy the condition of linear independence.

### 10 SAAF distribution: solution of one-dimensional diffusion equation and constant decay rate model

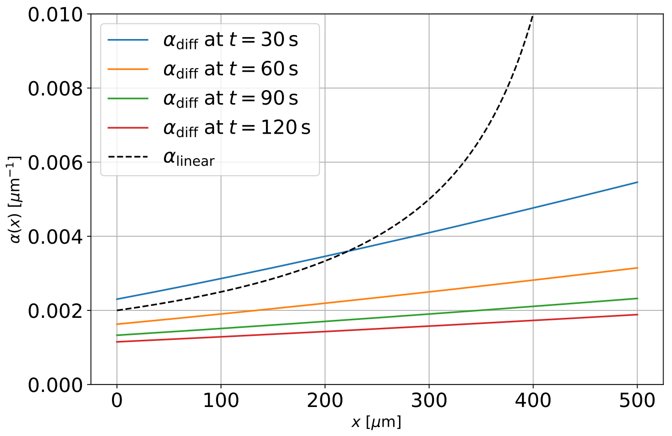


Figure 4: Spatial distribution of the decay rate for a solution of the diffusion equation and the linear distribution for $x_0 = 500$ [$\mu$m] and $D = 2000$ [$\mu$m$^2$/s]. For the solution of the diffusion equation, snapshots at $t = 30, 60, 90$ and 120 are shown. The dotted line at $\alpha = 0.002$, corresponding to Fig. 4, is also drawn.

In our experiment with a one-dimensional, linearly distributed SAAF field, we used a constant decay rate distribution to define a canonical trajectory for our model. Here, we compare this constant decay rate distribution with the solution of the one-dimensional diffusion equation—which provides a physically appropriate distribution of SAAF—to discuss the discrepancy between these models.

The one-dimensional diffusion equation with the Dirichlet boundary condition at $x = 0$ is defined as follows:

$$\frac{\partial s}{\partial t} = D\frac{\partial^2 s}{\partial x^2}, \quad (t > 0, x \in (0, \infty)), \quad s(t, 0) = c_0, \quad s(t, \infty) = 0,$$

where $D$ denotes the diffusion constant. This equation yields a self-similar solution:

$$s(t, x) = s_0 f(z) = s_0 \mathrm{Erfc}\left(\frac{z}{2\sqrt{D}}\right), \quad z = \frac{x}{t^{1/2}},$$

where $s_0$ is a constant. The decay rate of this solution, $\alpha_{\mathrm{diff}}$, is given by:

$$\alpha_{\mathrm{diff}} = -\frac{\partial s/\partial x}{s} = \frac{\frac{1}{\sqrt{\pi D t}}\exp\left(-\frac{z^2}{4D}\right)}{\mathrm{Erfc}\left(\frac{z}{2\sqrt{D}}\right)}.$$

The decay rate of the linear SAAF distribution, $\alpha_{\mathrm{linear}}$, is:

$$\alpha_{\mathrm{linear}} = \frac{1}{x - x_0}.$$

In Fig. 4, we plot $\alpha_{\mathrm{diff}}$ and $\alpha_{\mathrm{linear}}$ for $D = 2000$ [$\mu$m$^2$/s] and $x_0 = 500$[$\mu$m]. The distribution of $\alpha_{\mathrm{diff}}$ is not constant, but its variation is not significant, suggesting that our model exhibits similar behaviour to the case with a constant decay distribution of SAAF. Conversely, $\alpha_{\mathrm{linear}}$ has a stronger position dependency, which implies that a globally linear distribution of attractant does not provide position-independent trajectories in our model.
